# Functional mapping of the somatosensory cortex using noninvasive fMRI and touch in awake dogs

**DOI:** 10.1101/2023.12.22.572785

**Authors:** C.-N. Alexandrina Guran, Magdalena Boch, Ronald Sladky, Lucrezia Lonardo, Sabrina Karl, Ludwig Huber, Claus Lamm

## Abstract

Dogs are increasingly used as a model for neuroscience due to their ability to undergo functional MRI fully awake and unrestrained, after extensive behavioral training. Still, we know rather little about dogs’ basic functional neuroanatomy, including how basic perceptual and motor functions are localized in their brains. This is a major shortcoming in interpreting activations obtained in dog fMRI. The aim of this preregistered study was to localize areas associated with somatosensory processing. To this end, we touched N = 22 dogs undergoing fMRI scanning on their left and right flanks using a wooden rod. We identified activation in anatomically defined primary and secondary somatosensory areas (SI and SII), lateralized to the contralateral hemisphere depending on the side of touch, as well as activations, beyond an anatomical mask of SI and SII, in the cingulate cortex, right cerebellum and vermis, and the Sylvian gyri. These activations may partly relate to motor control (cerebellum, cingulate), but also potentially to higher-order cognitive processing of somatosensory stimuli (rostral Sylvian gyri), and the affective aspects of the stimulation (cingulate). We also found evidence for individual side biases in a vast majority of dogs in our sample, pointing at functional lateralization of somatosensory processing. These findings not only provide further evidence that fMRI is suited to localize neuro-cognitive processing in dogs in vivo, but also expand our understanding of touch processing in mammals, beyond classically defined primary and secondary somatosensory cortices.

**Significance Statement:** To understand brain function and evolution, it is necessary to look beyond the human lineage. This study provides insights into the engagement of brain areas related to somatosensation using whole-brain non-invasive neuroimaging of trained, non-sedated, and unrestrained pet dogs. It showcases again the usefulness of non-invasive methods, in particular fMRI, for investigating brain function and advances the mapping of brain functions in dogs; using this non-invasive approach without sedation, we are able to identify previously unknown potential higher-order processing areas and offer a quantification of touch processing lateralization.

## 1. Introduction

In recent years, canine neuroimaging has seen a rise in interest (Berns et al., 2012; Bunford et al., 2017; Huber & Lamm, 2017). The dog as a model organism provides us with interesting features, such as similar living conditions as humans (ManyDogs Project et al., 2023) and a remarkable social cognitive ability (Hare & Tomasello, 2005; ManyDogs Project et al., 2023; Topál et al., 2009).

To understand how higher order cognition is processed in the doǵs brain, we also need to understand the braińs fundamental organization and how it processes sensory input at the lower levels. To this end, recent research has mapped out the visual, olfactory and auditory cortices in awake and unrestrained dogs using functional magnetic resonance imaging (fMRI; Andics et al., 2014, 2016; Boch et al., 2021, 2023; Bunford et al., 2020; Cuaya et al., 2016, 2022; Dilks et al., 2015; Gillette et al., 2022; Jia et al., 2014; Phillips et al., 2022). However, our best understanding of the canine somatosensory cortex dates back to almost 70 years ago (Hamuy et al., 1956), using invasive methods, a small sample and focusing on selected parts of the canine cortex. Even in humans, invasive methods have limitations in the investigation of the somatosensory cortex (see Gordon et al., 2023). In this study, we examined somatosensory processing in a sample of healthy and awake pet dogs. Our main aims were to understand how a non-primate mammal processes touch, whether their touch processing is lateralized, and what parts of the brain may be involved beyond the primary and secondary somatosensory areas. With this approach, our study aimed to provide another important puzzle piece in comprehensively mapping and understanding dogs’ functional neuroanatomy in vivo and using non-invasive methodology.

During fMRI scanning, dogs were dynamically touched using a wooden rod moved down their left and right flanks. In line with our preregistration (https://osf.io/4gs9d/), we expected to see activation in the primary and secondary somatosensory cortices in response to touch (hypothesis 1), and that this activation would be higher in the hemisphere contralateral to the touched flank in primary somatosensory cortex SI, which in the dog is comprised of the postcruciate and rostral suprasylvian gyri, and in the ipsilateral hemisphere for secondary somatosensory cortex SII, the rostral ectosylvian gyri (Hamuy et al., 1956). Additionally, we hypothesized potential ipsilateral activation in the cerebellum, due to its (ipsilateral) involvement in motion control and potential involvement in suppressing touch elicited motion in our task (Uemura, 2015). Hypothesis 2 focused on whether somatotopic mapping could be achieved with the relatively coarse resolution of fMRI (Shmuel et al., 2007). In particular, we were interested in how activation shifts as a function of the dynamic somatosensory stimulation, possibly showing the trajectory of activation along the receptive fields coding for the parts of the back that were being stimulated, but also the progression of the signal into other parts of the canine cortex, to higher-order processing steps.

Moreover, we were interested in lateralization of somatosensory processing (hypothesis 3). While dogs may not possess a population wide side preference for one paw (Ocklenburg et al., 2019; Wells et al., 2018; but see Laverack et al., 2021), like humans favoring their right limbs (Papadatou-Pastou et al., 2020) or great apes do (Güntürkün et al., 2020; Hobaiter & Byrne, 2013; Hopkins, 2006), individually, most dogs seem to favor one over the other limb consistently (Ocklenburg et al., 2019). Even though somatosensation is not to be equated to motor behavior, we were interested in seeing whether somatosensation may be processed more strongly by one over another hemisphere within individuals (looking at laterality quotients, hypothesis 3). Based on the existing literature for motor biases (Charlton & Frasnelli, 2023; Ocklenburg et al., 2019; Simon et al., 2022), we expected the majority of dogs in our sample to show lateralized somatosensory processing.

## 2. Methods

### 2.1. Sample

Dogs in our sample had undergone extensive prior training to enable awake and unrestrained MRI scanning (Karl, Boch, Virányi, et al., 2020) and successfully participated in several prior studies (Boch et al., 2021, 2023; Guran et al., 2023; Karl, Boch, Zamansky, et al., 2020; Karl et al., 2021). For the present study, they additionally underwent familiarization with the touching rod and touching procedure. Dogs (and their owners) had been recruited through the Clever Dog Lab of the Messerli Research Institute at the University of Veterinary Medicine Vienna. Our final sample consisted of 22 dogs (9 female; mean age 4 years, age range: 1-8 years; mean weight = 21.9 kg). The sample consisted of a variety of fur types, as well as breeds, with 10 mixed-breed dogs, 5 retrievers, 6 shepherds, and 1 hunting dog. Our data collection stopping rule is specified in our analysis preregistration (https://osf.io/tdzrf), whereby we included all dogs that concluded two successful runs with less than 50% excluded volumes per run, by June 15^th^ 2023.

All dogs were examined for their generally good health condition and eyesight at the Small Animals Clinics of the University of Veterinary Medicine Vienna prior to inclusion into the dog imaging cohort. Owners gave written informed consent before data collection but no monetary compensation was given to dog owners for their dog’s participation. This work was approved by the institutional ethics and animal welfare commission in accordance with Good Scientific Practice (GSP) guidelines and national legislation at the University of Veterinary Medicine Vienna (ETK-06/06/2017), based on a pilot study conducted at the University of Vienna, and complies with the ARRIVE Guidelines (Kilkenny et al., 2010).

### 2.2. Touch stimulation procedure

Dogs lay on the scanner bed head first, in prone position, facing the back of the scanner. The dog trainers who performed the stimulation stood at the front of the scanner, to the right side of the scanner bed to touch the dogs with the rod. Trainers got auditory cues where to touch the dog, when to start, and when to stop. The rod was extended with another wooden stick to make sure that trainers did not need to reach into the scanner bore with their arms, potentially causing artefacts. We used a wooden backscratcher to touch the dogs on the right and left flanks, starting at the shoulder, all the way down to the hip joint (see Figure 1). One dynamic stimulation lasted 4 seconds, and was repeated once (with a 1s break for the trainer to reposition the rod at the shoulder), before a baseline block (10 s). Then, the other side was touched. In each run, 10 blocks of stimulations were administered, 5 on each side (alternating). Each dog in this sample successfully underwent two full touch runs. Touch stimulations were blocked (10 s) and alternated with 10 s baselines (no touch).

**Figure 1:**
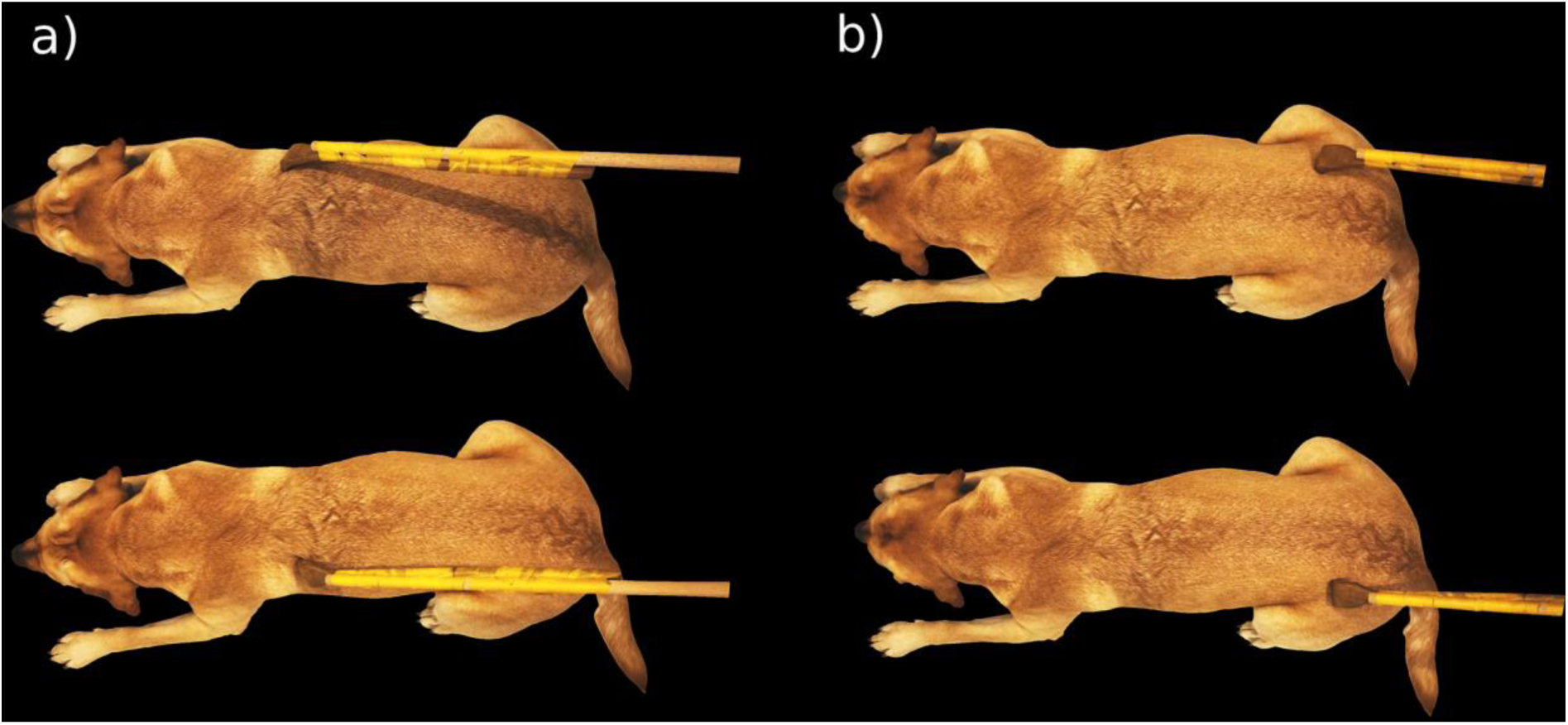
Illustration of the touching procedure and location. Touching started behind the left or right shoulder (a) and progressed for four seconds until reaching the hip area (b). Then, the other respective side was stimulated in the same way.

### 2.3. Data acquisition

Functional imaging data were obtained from 24 axial slices (interleaved acquisition in descending order, spanning the whole brain) using a twofold multiband-accelerated echo planar imaging (EPI) sequence with a voxel size of 1.5 x 1.5 x 2 mm^3^ (TR/TE = 1000/38 ms, FoV = 144 x 144 x 58 mm^3^, flip angle = 61°, 20% slice gap). The functional touch runs consisted of 215 volumes each. Structural images (134 volumes) were obtained using a voxel size of 0.7 mm isotropic [TR/TE = 2100/3.13 ms, field of view (FoV) = 230 x 230 x 165 mm^3^]. Runs could be acquired in the same or in separate sessions, depending on the dog’s capacity to lie still in the scanner for one or two runs in one session. We used a Siemens Magnetom Skyra (Siemens Healthcare, Erlangen, Germany) with a field strength of 3 Tesla for all measurements, and a head coil specifically designed for and tailored to optimal signal acquisition in dog fMRI (Guran et al., 2023).

### 2.4. Data preprocessing

Data preprocessing was performed using SPM12 within MATLAB version 2022a. Images were slice-time corrected to the middle slice and realigned. Thereafter, we performed manual reorientation for the structural and EPI images. Structural images were manually skull-stripped with itk-SNAP (Yushkevich et al., 2006). This step is of particular importance in dog MRI, where the skull is bordered by massive musculature which can hinder successful coregistration, which was performed onto the mean image of each run. Structural segmentation of the brain was performed using the canine tissue probability maps provided by Nitzsche et al. (2019). Normalization of functional and structural data were performed using the “Old Normalization” module in SPM (originally implemented in SPM8), finally reslicing images to 1.5-mm isotropic voxel size, and smoothing of 3 mm (with a Gaussian FWHM kernel). Data were motion scrubbed by calculating framewise displacement, and excluding volumes with a displacement larger than 0.5 mm in comparison to the previous volume (Power et al., 2012, 2014). In our final sample, we excluded an average of 15.3% of volumes in each run (roughly 33 volumes). Note that as part of our overall quality assurance measures, runs with ≥ 50% of volumes exceeding this threshold were discarded, and repeated in another imagining session (until this criterion was fulfilled for two total runs per dog).

### 2.5. Analyses

We used different analysis approaches based on our questions of interest. To localize somatosensory activation, we used mass-univariate whole-brain and regions of interest GLM analyses in SPM12 using a dog-tailored canonical hemodynamic response function (Boch et al., 2021). To test for somatotopic activation shifts as a function of time, we used a time-resolved finite impulse response (FIR) analysis.

#### 2.5.1. GLM analysis

We modelled the events of interest using a design matrix with the following regressors: Touch Left and Touch Right. From this we defined the following contrasts: Touch vs Baseline (T vs BL), which contrasted activation in response to left and right-sided touch to the baseline (no touch blocks, visual fixation), Right touch vs Baseline (R vs BL), where the right flank of the dog was touched, Left touch vs Baseline (L vs BL), where the left flank was touched. These contrasts were defined on a subject level (1^st^ level) and then entered into a second level GLM analysis. In addition to a whole-brain exploration of activation in these contrasts, we tested *a priori* hypotheses on somatosensory engagement with higher sensitivity. To this end, we defined anatomical regions of interest (ROIs) for primary and secondary somatosensory cortex SI and SII, and used these as masks for an ROI analysis on the 2^nd^ level, using the small volume family-wise error correction utility in SPM12. Specifically, ROIs were based on previous invasive literature (Hamuy et al., 1956) and included the postcruciate and the rostral suprasylvian gyri as SI, and the rostral ectosylvian gyrus as SII, see Figure 2.

**Figure 2:**
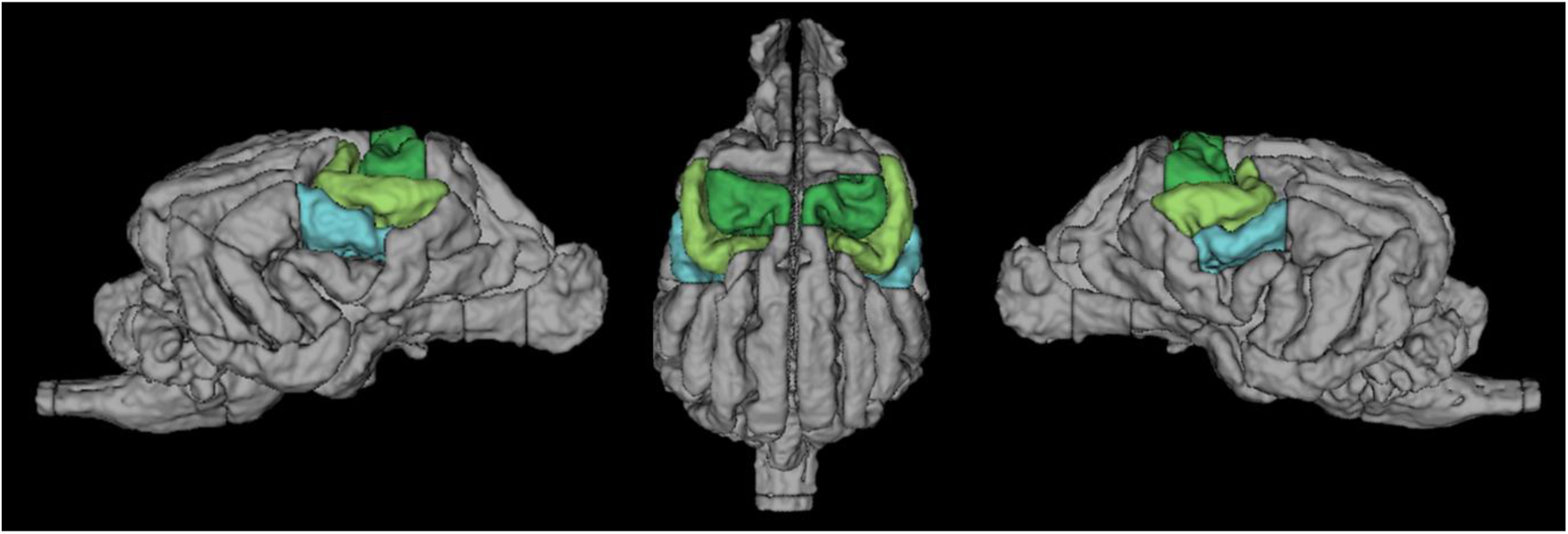
Location of SI in green, SII in light blue. Dark green = Postcruciate gyri. Light green = rostral suprasylvian gyri. Light Blue = rostral ectosylvian gyri (Czeibert et al., 2019).

Anatomical regions underlying clusters outside of the anatomical SI+SII mask refer to a dog anatomical atlas (Czeibert et al., 2019) which was normalized to a breed-averaged template space (Nitzsche et al., 2019). For statistically thresholding the whole-brain analyses, we had preregistered based on prior research a cluster-level inference approach with a cluster-defining threshold of p < 0.005 (FWE-corrected at cluster level, α = 0.05); since the resulting clusters were unexpectedly large, we also used these criteria for the small-volume-correction analyses; note that this is a deviation from the preregistration (where we had preregistered voxel-level FWE-correction of α = 0.05).

##### 2.5.1.1. FIR analysis

To ascertain whether “traveling” of activation across the cortex, in line with “caninculi” (in analogy to the creatures of the human somatosensory cortex, the homunculi) found in previous research (Penfield & Jasper, 1954), or potential higher order processing, can be seen using fMRI while stimulating in a blocked design, we used a FIR analysis approach. We used bins of 1s each, thus splitting each stimulation (two per block) into four bins, and to visualize potential travel of activation across the stimulation. This analysis was performed separately for left and right touch stimulation. We compared the 1^st^ and 4^th^ (last) bin using T-tests. This contrast was chosen to maximize power for finding activation location differences. However, since we discovered no significant clusters in the fourth time bin, we also compared the first and third time bins. Finally, we compared each time bin to baseline levels and also inspected activation shifts across the time bins visually.

#### 2.5.2. Lateralization analysis

To investigate lateralization of activation, we extracted beta values from right and left hemispheric anatomically defined unthresholded clusters in the T vs BL contrast. We computed laterality quotients of beta weights per cluster, using this formula:

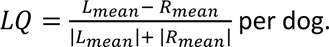

Then, we used the absolute values of all dogs to run a t-test comparison against 0, to ascertain whether the sample showed lateralization (regardless of direction). We only considered dogs as lateralized if they had an LQ over 0.1 or under −0.1 (a more conservative threshold than used elsewhere, see Zahnert et al., 2022; for a discussion of LQs see Seghier, 2008), took the absolute of the bias and ran a t-test with them, testing against a 0 mean (no lateralization).

### 2.6. Open science, data and code availability

We follow open science practices throughout, including pre-registration of the analyses, which can be found here: https://osf.io/tdzrf; canine fMRI data (2^nd^ Level contrast SPM maps), as well as analysis scripts are available on github (https://github.com/alexandrinaguran/somatostudy).

## 3. Results

### 3.1. GLM results

#### 3.1.1. Touch vs Baseline

To understand touch processing in the dog brain, we first ran t-contrasts in SPM using an anatomical mask of the SI and SII as an inclusive mask. Contrasting touch (bilateral) vs baseline, we found bilateral clusters located in the somatosensory cortex of the dogs, in a left hemispheric cluster (peak location: −20 −4 12) with k = 65, p_FWE_ < 0.001, T = 4.98, a right hemispheric cluster (peak location: 20 −6 12) with k = 23, p_FWE_ < 0.05, T = 5.65, and a medial cluster (only at trend level, peak location: 2 0 19, k = 21, p_FWE_ = 0.053, T = 4.53), see Figure 3A. There is a stronger activation in the touch condition, as opposed to the baseline condition. In the reverse contrasts (baseline > touch), investigating relative deactivation during the touch vs. baseline, we found two clusters, one right hemispheric (peak location: 12 2 19) with 105 voxels, p_FWE_ < 0.001, T = 5.26, and one left hemispheric (peak location: −16 2 14) with 63 voxels, p_FWE_ < 0.005, T = 5.05, see Figure 3B (i.e. higher activation in the Baseline condition than during touch).

**Figure 3:**
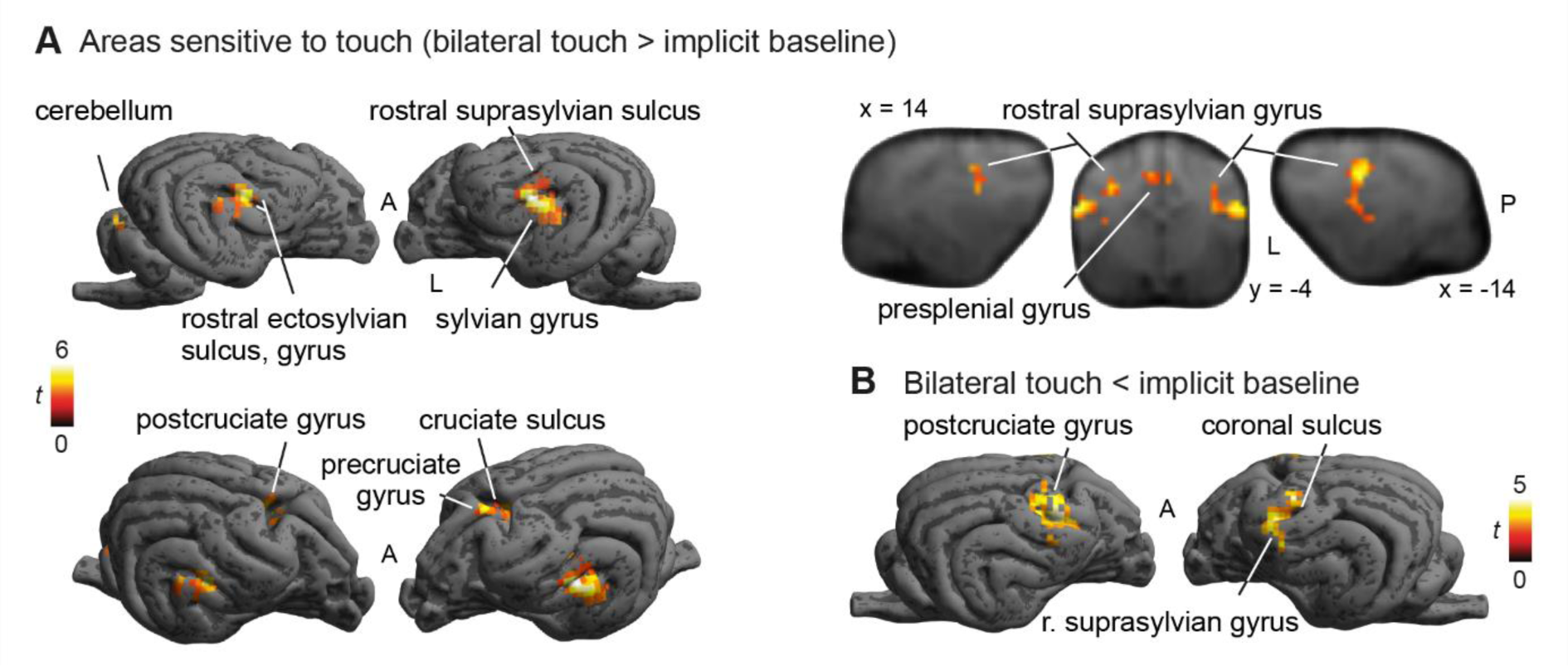
A) Activations in response to Touch stimulation (whole-brain), N = 22. Bilateral SI and SII activation, as well as medial cluster (whole-brain), presumably in the cruciate sulcus and precruciate, as well as cingulate, gyrus. B) Stronger responses in baseline than touch predominantly in postcruciate gyri. Image cluster thresholded at .005, 40 voxels.

On the whole-brain level (without any anatomical or any other mask), all three cortical clusters reached highly significant levels (all p_FWE_ < 0.005), for both activations and deactivations. However, the clusters with stronger activation during touch vs baseline showed strongly increased extent in the whole-brain analysis, e.g. more than twice the size for the first cluster, and even larger differences for the other clusters (see Table 1), and were located more posteriorly than the clusters in the reverse contrast (see Figure 3 B). In addition to the cortical clusters, we found a cerebellar cluster (peak location: 5 −39 4) with k = 42, p_FWE_ = 0.038, T = 4.81, with stronger activation during touch vs baseline.

**Table 1:**
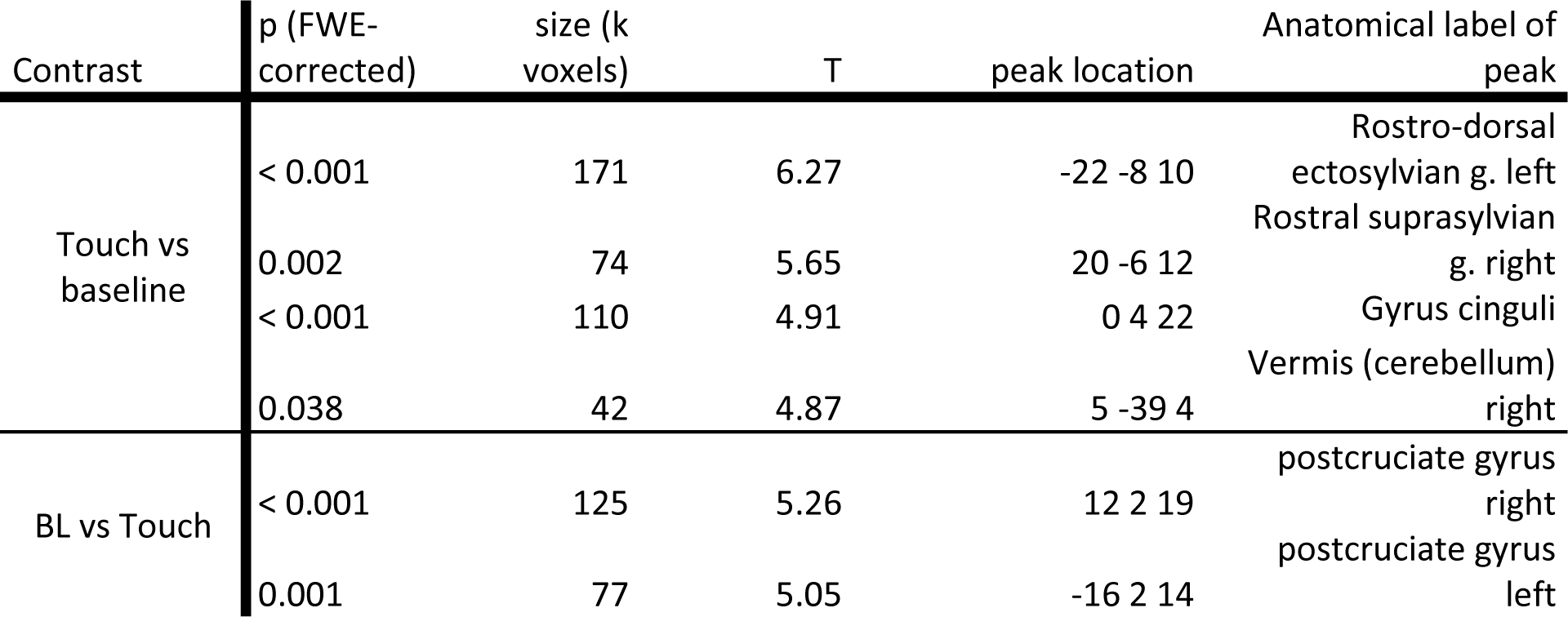
Significant clusters in the wholebrain analysis, contrasting Touch (both sides) with Baseline. Clusters are similar to those found in the svc (see peak locations), however much larger and have generally higher T-values and better significance.

#### 3.1.2. Left-sided touch vs Baseline

Contrasting touch on the left flank vs baseline, we found a large cluster in the right hemisphere (peak location: 20 −6 12) with k = 74, p_FWE_ < 0.001, T = 6.9, as well as a smaller left hemispheric cluster, which only reached trend level in the SVC analysis (peak location: −16 −8 12) with k = 18, p_FWE_ = 0.073 (cluster level), T = 4.75, and a central cluster (peak location: 5 0 19) with k = 30, p_FWE_ = 0.016, T = 4.68, which showed stronger activation during left flank touch than baseline. In the opposite direction, we found one right hemispheric cluster (peak location: 14 4 20) with k = 70, p_FWE_ < 0.001, T = 6.18, showing stronger activation in Baseline than touch, see Figure 4).

**Figure 4:**
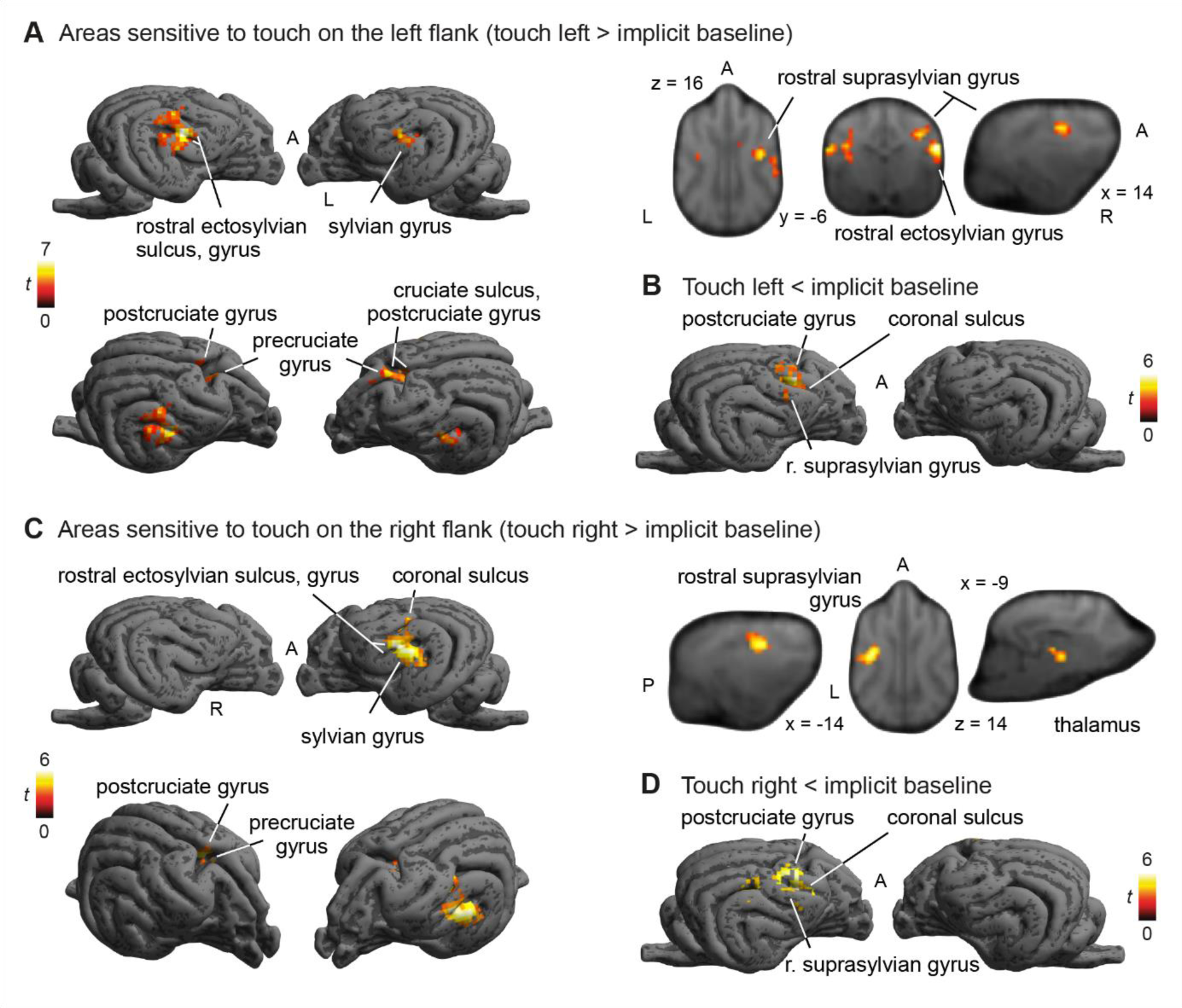
BOLD response to touch on left flank (A+B) and right flank separately (C+D). Strong activation in posterior parts of SI and SII and rostro-medial regions. De-activation in postcruciate gyrus (B +D). Image cluster thresholded at .005, 50 voxels.

On the whole-brain level, a similar picture as in the touch vs BL analysis arose: descriptively all clusters were larger with smaller p-values (see Table 2).

**Table 2:**
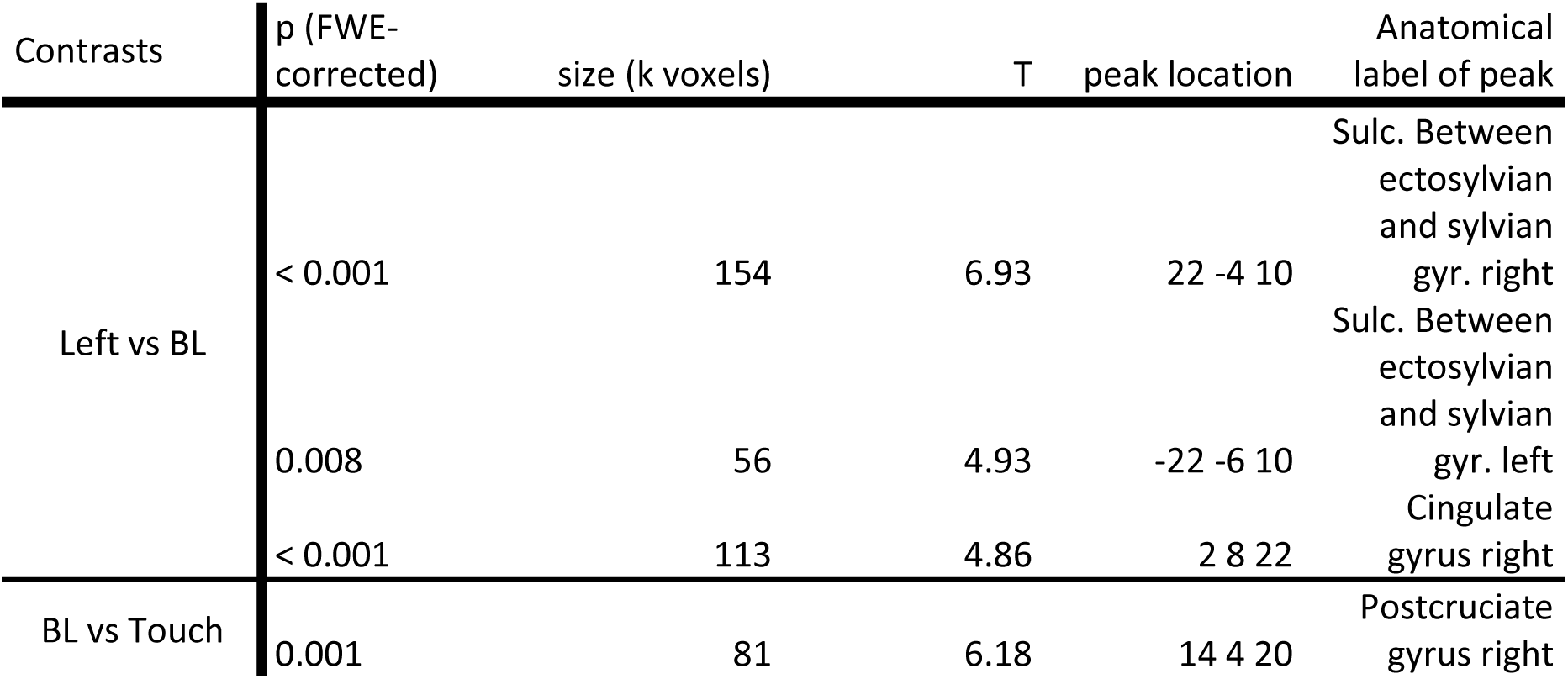
Significant clusters in the whole-brain analysis of left flank touch vs baseline.

| Contrasts | p (FWE-corrected) | size (k voxels) | T | peak location | Anatomical label of peak |
| --- | --- | --- | --- | --- | --- |
| Left vs BL | < 0.001 | 154 | 6.93 | 22 -4 10 | Sulc. Between ectosylvian and sylvian gyr. right |
|  | 0.008 | 56 | 4.93 | -22 -6 10 | Sulc. Between ectosylvian and sylvian gyr. left |
|  | < 0.001 | 113 | 4.86 | 2 8 22 | Cingulate gyrus right |
| BL vs Touch | 0.001 | 81 | 6.18 | 14 4 20 | Postcruciate gyrus right |

#### 3.1.3. Right vs Baseline

Contrasting touch on the right flank with baseline, we found a large cluster in the left hemisphere (peak location: −19 −4 12) with k = 83, p_FWE_ < 0.001, T = 5.73, with stronger activation for touch on the right flank vs baseline (see Figure 4C). In the opposite direction, we found one right hemispheric cluster (peak location: 12 2 18) with k = 79, p_FWE_ < 0.001, T = 5.97, with stronger deactivation during right flank touch, see Figure 4D.

Again, on the whole-brain level, we replicated all clusters we found in the SVC analysis, with larger cluster sizes. Additionally, we found a central cluster as well as a left hemispheric cluster (see Table 3).

**Table 3:**
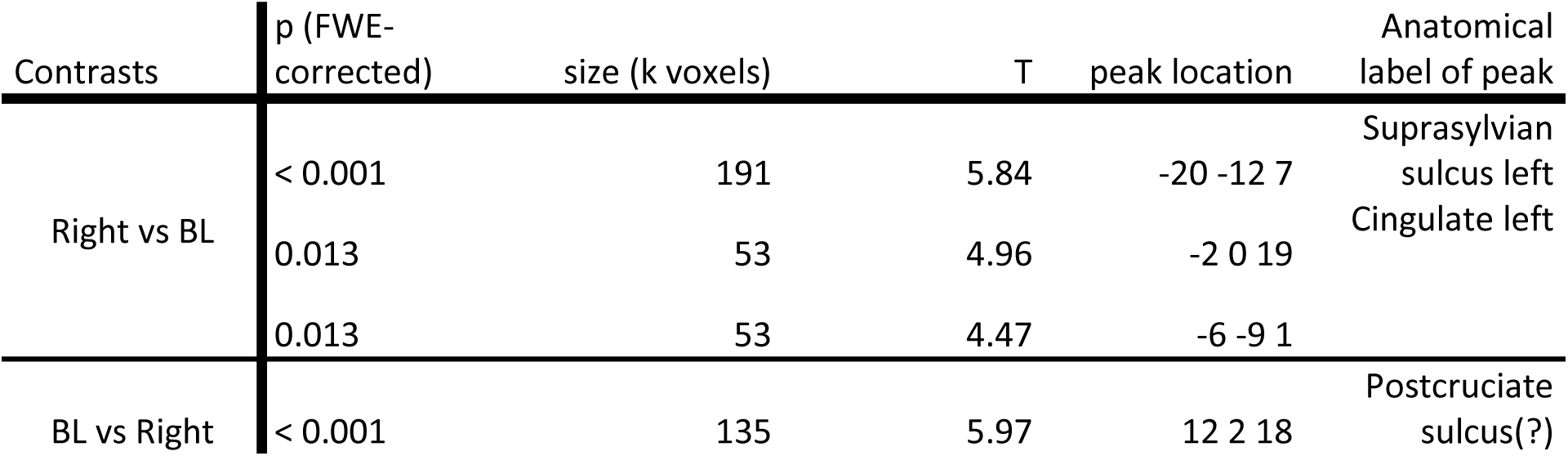
Significant Clusters in the whole-brain analysis, contrasting Right touch and Baseline. Note the additional central and left hemispheric clusters, absent in the SVC analysis.

#### 3.1.4. Investigating activation outside of anatomical masks

Through the GLM analysis and the notably larger clusters in the whole-brain analysis, it is apparent that the overlap between the observed activation and the regions we expected to be activated (as included in our anatomical masks) was not optimal. Therefore, we ran an exploratory svc analysis using our anatomical mask as an *exclusive* mask, purely to better understand and illustrate the location of the activation outside of the anatomical mask. We found 4 significant clusters, one left hemispheric cluster, located close under SII, p_FWE_ = 0.001, k = 84, T = 6.27, peak location at −22 −8 10, one right hemispheric cluster, p_FWE_ = 0.038, k = 42, T = 5.16, peak location at 20 −6 10. Additionally, we found one rostro-central cluster, p_FWE_ < 0.005, k = 73, T = 4.91, peak location at 0 4 22, and a cerebellar cluster, p_FWE_ = 0.038, k = 42, T = 4.87, at 5 −39 4 (all p from cluster level inference). There were no significant clusters in the reverse contrast (BL vs Touch). The left and right hemispheric clusters were located in the rostral and caudal sylvian gyri (left and right respectively), the central cluster comprised parts of the right cingulate gyrus and right precruciate gyrus. The cerebellar cluster was located in the right hemisphere and parts of the (right) vermis. Figure 5 shows all activations and deactivations in the main contrast (Touch vs BL) with the contours of the anatomical mask superimposed. Please note that this analysis was not preregistered. We used the same inference levels as for all other analyses.

**Figure 5:**
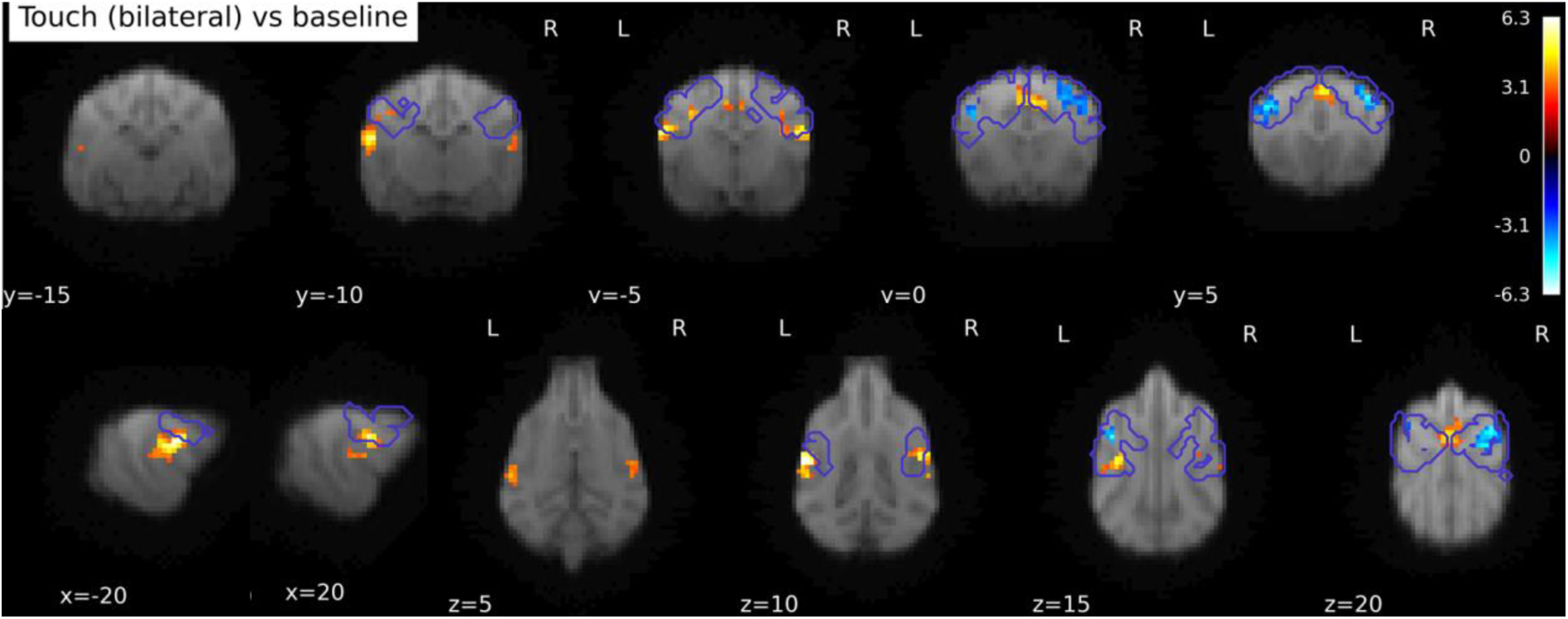
Activation in Touch vs Baseline contrast, whole-brain. Purple contours: somatosensory cortex mask bounds (SI + SII). Clusters outside of SI+SII are ventral and caudal, as well as central. Image cluster thresholded at .005, 40 voxels.

### 3.2. FIR analysis

#### 3.2.1. First vs Last time bins

As above, we used an anatomical mask of SI and SII for SVC in these analyses, when not specified differently. Contrasting the first second of touch stimulation on the right flank with the last (4^th^) second of right touch stimulation, we found only one cluster at trend level, p_FWE_ = 0.071, k = 35, T = 4.02, with peak location at −16 −6 18. This activation is significant when exploring the data using a mask that only included the contralateral left hemispheric SI and SII, with p_FWE_ < 0.005, see Figure 6. There were no significant clusters in the opposite direction (4^th^ time bin vs first time bin). For the touch stimulation of the left flank, there was again a trend level cluster, p_FWE_ = 0.078, k = 17, T = 3.89, with the peak location at the exact contralateral side as found in the right flank simulation, at 16 −6 18. Again, using small-volume correction for the right hemispheric SI + SII, this cluster reached significance, p_FWE_ < 0.005. There were no significant clusters in the opposite direction (last time bin vs first time bin). None of the clusters survived on the whole-brain level.

**Figure 6:**
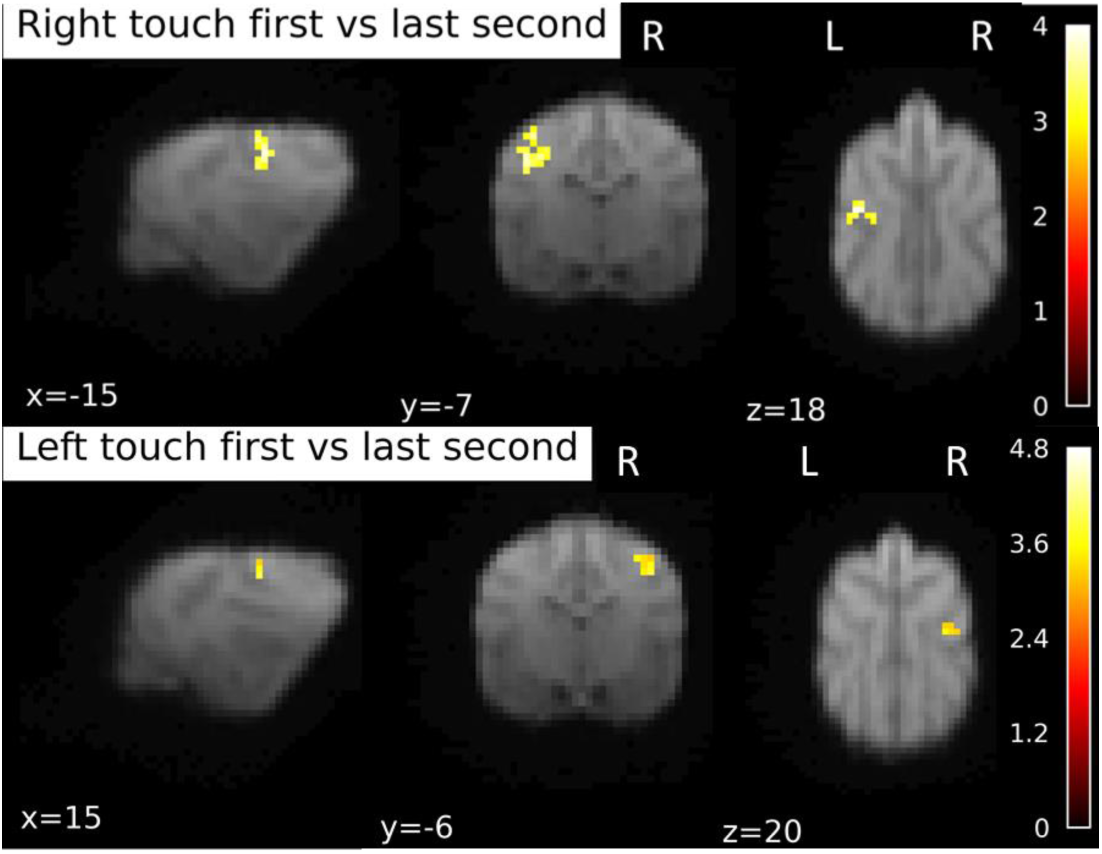
BOLD response during first second of touch stimulation contrasted to the last second of stimulation. Separately for right and left flank touch. Image of right touch cluster thresholded at .1 (to capture the trend level activations), 15 voxels. For left touch, .05 and 10 voxels.

#### 3.2.2. Time bins vs Baseline

In addition to contrasting the two time bins where we expected the strongest difference (first and last), we looked at activation differences against baseline for each time bin, on a whole-brain level. In particular, we were interested in seeing whether the activation visibly shifts across the somatosensory cortex as time progresses. For the first time bin (first second of stimulation), we found four clusters for the Left touch (two in the right hemisphere, one medial, one in the left hemisphere, see Table 4. All clusters survived the whole-brain level. For the first second of right touch stimulation we found a large left-hemispheric cluster (k = 131), which not only survived on the whole-brain level, but was notably larger (k = 356), see Table 4. For the second and third time bins, there was at minimum one cluster in the hemisphere contralateral to the touch stimulation, which survived whole-brain level, see Table 5 and Table 6. There were no significant clusters in the fourth (last) time bin.

**Table 4:**
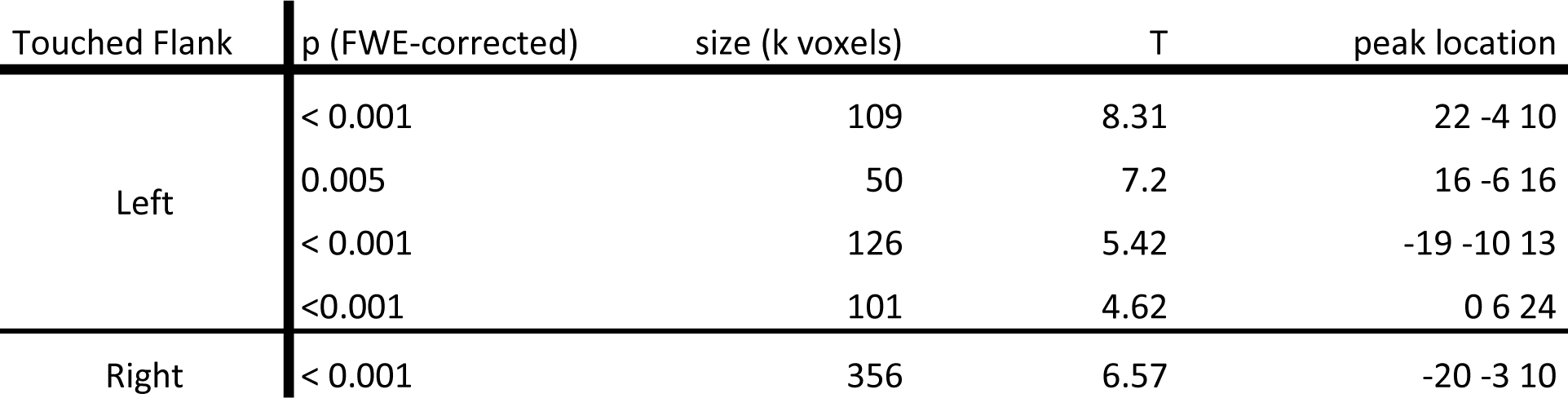
Significant clusters contrasting the first time bin against Baseline on a whole-brain level.

**Table 5:** Significant clusters contrasting the second time bin against Baseline on a whole-brain level.

| Touched flank | p (FWE-corrected) | size (k voxels) | T | peak location |
| --- | --- | --- | --- | --- |
| Left | 0.001 | 78 | 6.13 | 18 -4 12 |
| Right | 0.001 | 71 | 5.42 | 2 -2 16 |
|  | < 0.001 | 98 | 5.2 | -19 -9 16 |
|  | 0.043 | 38 | 3.75 | 2 -34 2 |

**Table 6:** Significant clusters contrasting the third time bin against Baseline on a whole-brain level.

| Touched Flank | p (FWE-corrected) | size (k voxels) | T | peak location |
| --- | --- | --- | --- | --- |
| Left | < 0.001 | 96 | 5.97 | 2 0 19 |
| Right | 0.008 | 53 | 6.34 | -13 -4 16 |
|  | 0.001 | 72 | 3.99 | -1 0 19 |

##### 3.2.2.1. Visual Inspection of activation shift

While statistical differences between time bins were observed, we also wanted to get an idea of the activation changes as a function of time. To this end we inspected data visually, by plotting the significant clusters found in each time bin onto the Touch vs BL activation, see Figure 7 below. From the visual inspections, the following qualitative observations seem to be possible: 1) activation is particularly strong in the first bin, and virtually nonexistent in the final bin. 2) Activation shifts to more central areas from the first to the third bin. 3) The peak of cerebellar activity happens during the second time bin. 4) Activations in the second and third time bin seem more concentrated on central and dorsal areas, compared to activation during the first bin.

**Figure 7:**
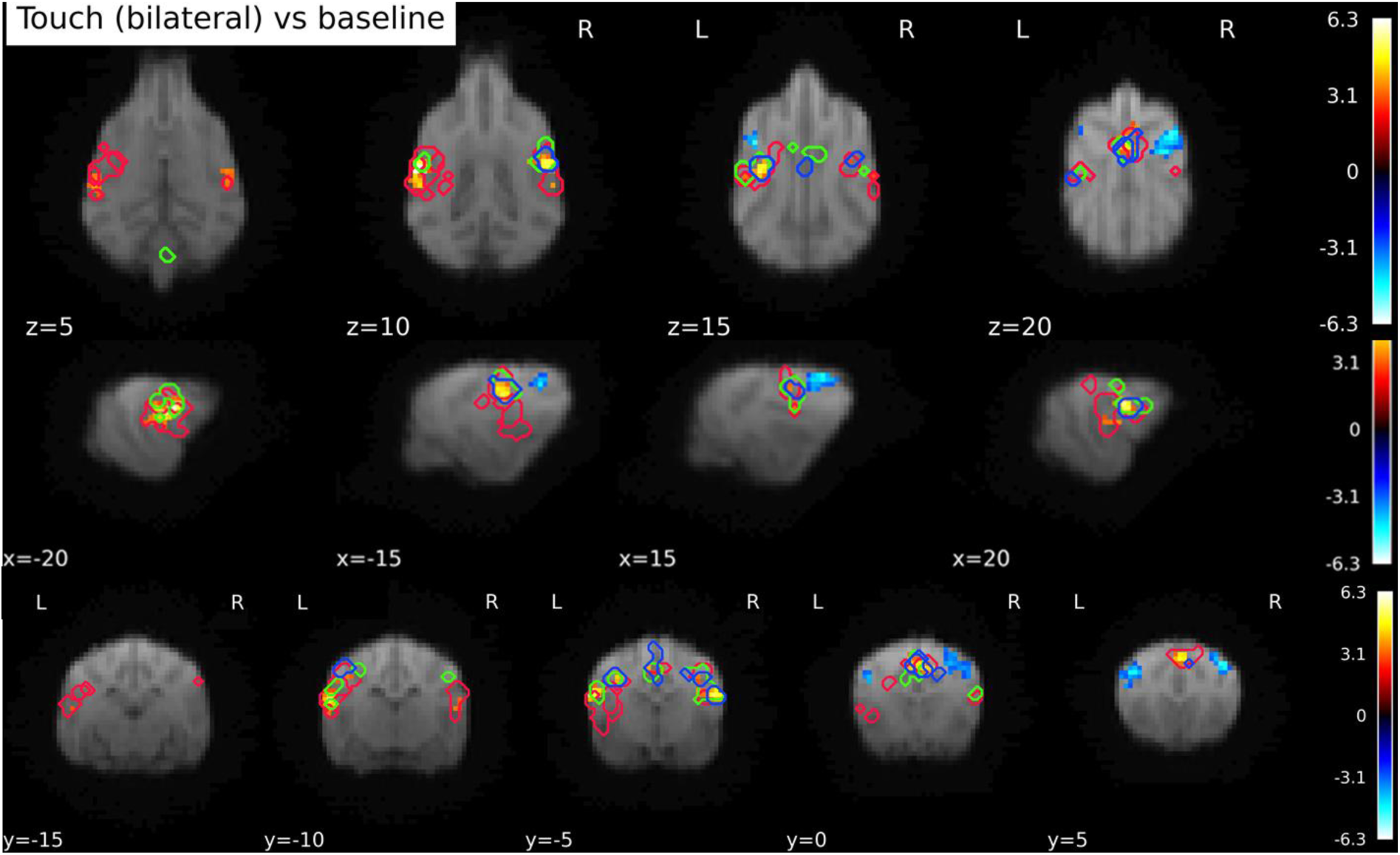
Clusters of activation by time bin, plotted on top of the Touch vs BL contrast. Red outlines mark the activation in response to touch in the first time bin (0-1s). Green outlines activation in the second time bin (1-2s), and blue the activation of the third time bin. No significant clusters were found in the fourth time bin. The figure illustrates both a relative overlap of activations, but also a shift towards more dorsal parts of SI and SII over the course of time.

#### 3.2.3. Comparison of first and third time bin

Since there were no significantly activated clusters in the 4^th^ time bin, we decided to run a comparison between the first and the third bin. Please note that this analysis was not preregistered or planned, but arose from the analysis of the data, i.e. the result that no significant clusters were found in the 4^th^ time bin.

Comparing the first and third time bin on the left and right hemisphere separately, again using SVC with an anatomical mask of SI and SII, we found a left hemispheric cluster, p_FWE_ < 0.001, k = 94, T = 6.12 with peak at −19 −4 12 in response to right touch, and a right hemispheric cluster, p_FWE_ < 0.001, k = 67, T = 7.63 with peak location at 20 −4 12, and a rostro-central cluster, p_FWE_ = 0.02, k = 23, T = 4.75, peak location at 2 0 19, in response to left flank touch, see Figure _8_.

**Figure 8:**
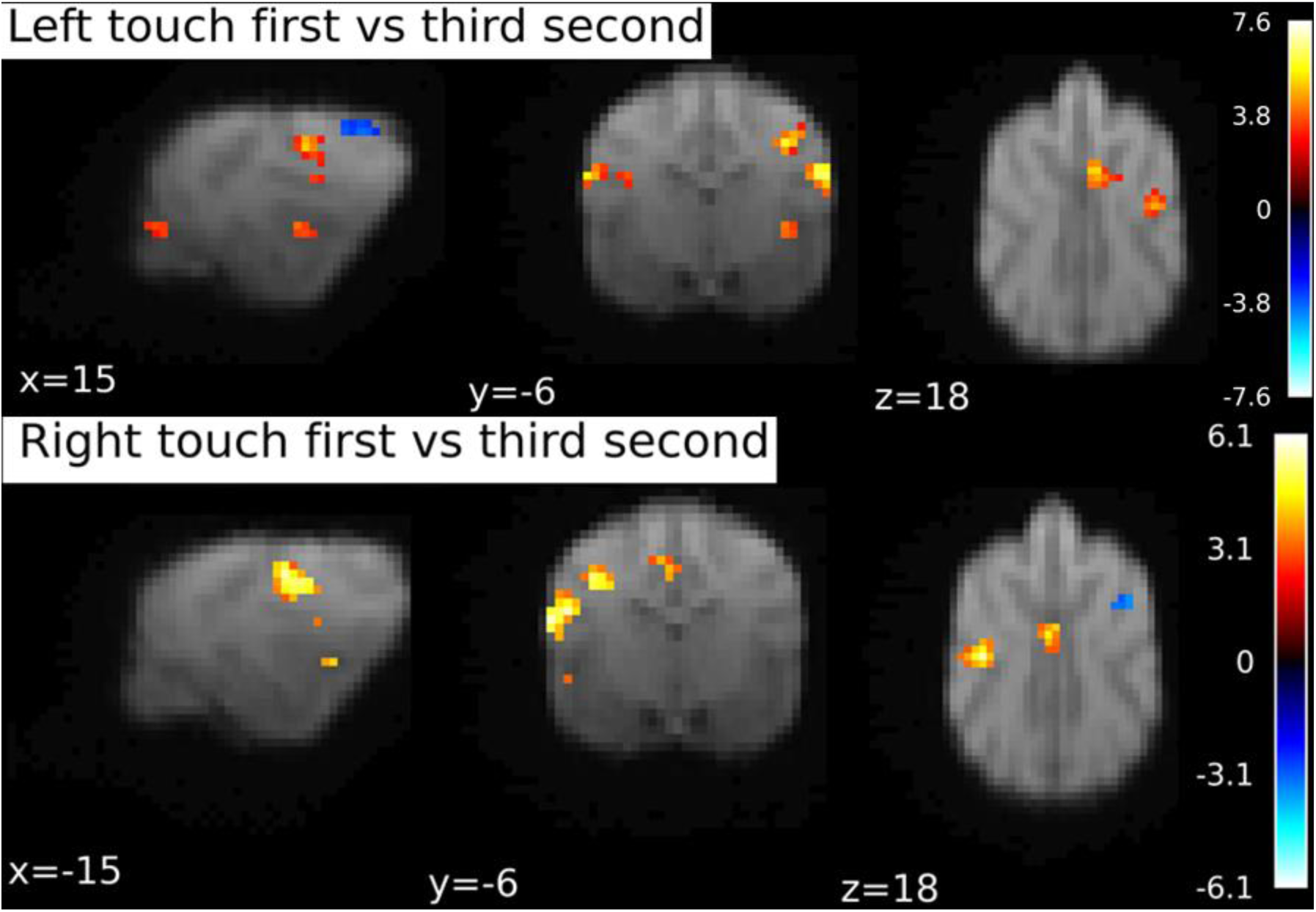
BOLD response during first second of touch stimulation contrasted to the third stimulation second. Separately for right and left flank touch. Image cluster thresholded at .05, 10 voxels.

### 3.3. Laterality quotients

Out of the sample of 22 dogs, 19 showed a lateralization, 9 towards the right hemisphere and 10 towards the left hemisphere (values smaller than −0.1 or bigger than 0.1 respectively). The mean absolute lateralization quotient was 0.49. For the dogs whose touch processing was lateralized to the left hemisphere, the mean was 0.54 (± 0.36 std), and for the right-hemisphere-processing dogs 0.58 (±0.4 std).

The sample showed a sample wide asymmetry, p_FWE_ < 0.001, T_21_ = 5.9 (undirected). This was also true for the left (p < 0.001, T_9_ = 4.71) and right (p_FWE_ < 0.005, T_8_ = −4.32) lateralized dogs separately.

Since there is some debate as to how to interpret negative mean beta weights, we plotted the left and right hemispheric cluster means per dog, see Figure 9 including areas where LQ quotients above 0.1 and below −0.1 can be found.

**Figure 9:**
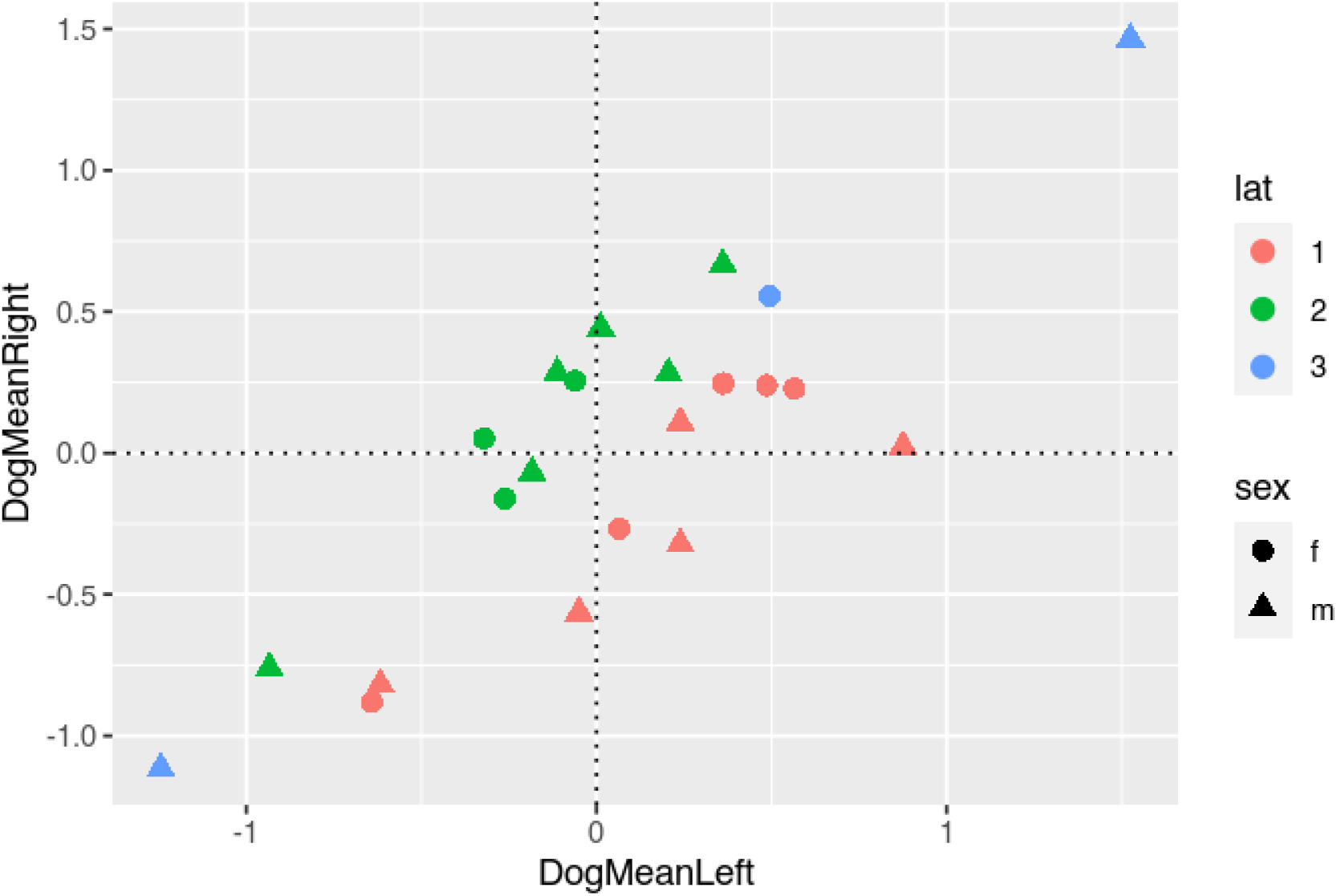
Relationship between mean left and right activation and final laterality indices (with absolute values in the denominator), see 2.5.2. Circles represent female, triangles male dogs. Blue markers indicate no lateralization (values between −0.1 and 0.1), green a rightward processing preference (LQ higher than 0.1), and red a leftward processing preference (lower than −0.1).

## 4. Discussion

To understand touch processing in awake, healthy dogs, we touched dogs on their right and left flanks while undergoing non-invasive brain scanning using fMRI. Building on invasive work from the 1950s, we found activation (and de-activation) in areas previously identified as SI and SII of the dog, but also activation in other brain areas. We also found evidence for lateralization of responses and a shift of functional activity hotspots in line with the dynamics of stimulation.

Firstly, we found clusters in the left and right hemisphere that responded more strongly to touch than baseline (no touch), as well as in a medial cluster. However, in addition to the hypothesized activation, we found additional areas outside the somatosensory areas that responded to touch stimulation. Touch activated the rostral and caudal sylvian gyri left and right, areas ventral and slightly caudal to SII, as well as the right cingulate cortex and right precruciate. The rostral sylvian gyrus has been linked to emotion perception (Hernandez et al. 2017; Karl et al., 2021), as well as the detection of familiar vs unfamiliar speech (Cuaya et al., 2022), and therefore seems to act as a multisensory area that could play a role in sensory integration and association processes. Interpreting the findings, it seems unlikely that the areas outside of SI and SII would be directly related to primary sensory processing, based on findings in dogs but also other mammalian species, when it comes to touch processing. Rather, they might be related to higher-order processing of touch. In the case of the gyrus cinguli, affective processing of the touch stimulation may be localized there, as the gyrus cinguli has been shown to relate to affective and even social processes in both humans and non-human primates (Devinsky et al., 1995; Rudebeck et al., 2006; Vogt et al., 2005) but also to motor control (Wang et al., 2001). Little is known about the functionality of the canine cingulate cortex so far, but it may play a role in integration of information and functional processing (Szabó et al., 2023). Additionally, the central cluster was biased towards the right side. In dogs, different side biases for processing of sensory inputs exist, and namely, the right hemisphere may play a larger role in processing arousing (or negative) stimuli (Simon et al., 2022; Siniscalchi et al., 2017). Through the domestication process and intensive handling by humans, experienced by most dogs, it is plausible that their brains show particular adaptations to process touch administered by humans in a social way, leading to increased arousal. Finding involvement of higher order cognitive processes highlights the methodological advantage of using a non-invasive method in healthy functioning animals: these findings can be made impossible when other parts of the brain are removed or damaged surgically (Hamuy et al., 1956; Marshall et al., 1937).

When contrasting left or right flank touch separately to baseline, we did find major activation in the contralateral hemispheres in locations consistent with SI/SII (rostral suprasylvian and ectosylvian gyri right, left suprasylvian gyrus). Ipsilateral processing was only found for the left touch, and also only at trend level in the SVC analysis, but remained absent for right flank touch. Peak locations of activations were almost identical for both hemispheres. These results suggest a strong preference for contralateral processing and potentially a bigger role of SI in flank stimulation, in line with previous findings on touch processing in mammals.

We also found cerebellar activation, which is generally associated with motor control and conditioning (Glickstein & Yeo, 1990; Ohyama et al., 2003): both aspects may have been relevant here, as the dogs are both relying on their learned behavior to lie still in the scanner and not respond with an orienting response to the touch stimulation, but also need to actively suppress motor responses.. This is also in line with the observed deactivation in the postcruciate gyrus (rostral SI right). Interestingly, we found the deactivation only in the right hemisphere, for both directions of touch. Due to its proximity to areas that have been suggested to play a role in motion (precruciate gyrus), it may be that the right postcruciate gyrus stands in close connection to downregulation of precruciate activity, playing a role in the inhibition of motor responses.

Our second hypothesis related to the possibility of tracking activation shifts across the 4s stimulation period starting at the shoulder and ending at the hip. Contrasting the first and third time bin, we found activation in the left and right hemispheres as well as a rostro-central cluster, aligning with parts of the postcruciate gyrus left and right: Importantly with time, processing moved away from the primary sensory areas and towards potentially higher-order association related parts of the cortex (see Figure 7), as revealed by higher engagement of rostro-central cortical areas. Thus, the FIR analysis approach allows some insights into the progression of sensory signal processing across higher-order cortical areas (i.e., outside the primary and secondary somatosensory cortices).

Finally, our third aim was to investigate whether dogs show lateralization (hemispheric dominance) of somatosensory processing. Already in the GLM analysis, we found some effects that indicated substantial lateralization, such as the deactivation in the right postcruciate gyrus. While somatosensory processing should not be equated with lateralized motor preferences, such as pawedness, which are more commonly investigated, lateralization in somatosensory processing can be related to motor biases and could be indicative of a generalized feature shared by many vertebrates (Güntürkün et al., 2020; Rogers et al., 2013). Roughly 86% of dogs (19 out of 22) showed a lateralization bias towards one hemisphere, as quantified through beta weights, a significant deviation from 0, with roughly equal numbers for left and right hemispheric biases. While there is no population wide preference for use of the left or right paw, around 70% of dogs generally show an individual side preference. This number is lower, but in a similar ballpark, than the number of dogs that showed a lateralized processing in our sample here. Thus, sensory processing is lateralized in most dogs. Thesedata give no reason to assume a population wide side bias of processing, in contrast to a leftward grey matter volume bias in dogs in general (Barton et al., 2023), suggesting a complex relation between functional and structural asymmetries.

Canine fMRI research is often limited by its sample sizes, an issue we circumvented by extending data collection until reaching a sample of 22 dogs, however, the future of canine and comparative neuroscience should be collaborative, to maximize power for important questions on canine cognition and cognitive evolution (ManyDogs Project et al., 2023).

In our approach of touch stimulation, a few limitations should be noted: first of all, the stimulations were performed by the dog trainers, and not a robotic apparatus. While timings and the procedure itself were both practiced and cued, it is very likely that a degree of variability in stimulations was present, which is of relevance to the FIR analysis, in addition to the coarse neural somatosensory representation of the back in comparison to other areas (Coq et al., 2004; Kaas, 1993; Woolsey et al., 1942). We chose this area pragmatically: stimulating the dog from the front would have brought the trainer into the view of the dog, possibly inducing distraction and other effects. From the back of the scanner, only the flanks and the paws were accessible, and many dogs do not like being touched on their paws and show this by moving them out of reach. Thus, only the back stimulation was viable. However, even though the back may not be the somatosensorily best represented area of the doǵs body, we were still able to find differences between touch bins on this area.

Finally, for the analysis of lateralization, within fMRI research, it is a continuing debate how to proceed with negative values. Since no consensus exists in the literature, one approach is to a) look at negative and positive values separately, or b) use absolute values in the denominator of the fraction used to calculate laterality indeces (Seghier, 2008). We opted for the latter version, since we would have shrunk the sample if only looking at either negative or positive values (not all dogs *had* both positive and negative values).

Using fMRI, we confirmed the roles of the dog’s postcruciate and rostral suprasylvian gyri (somatosensory area I) as well as the rostral ectosylvian gyri (SII) in the processing of touch. Beyond these areas that are likely most closely tied to sensory-perceptual computation, we pinpointed additional areas, namely parts of the rostral cingulate cortex, which may relate to affective processing, as well as the Sylvian gyri, which may play a putative role in higher-order or associative somatosensory processing. We found somatosensory processing to be lateralized in most dogs, with the sample being evenly split between left-, and right-hemisphere biased dogs. First indications for a somatotopic organization based on analyses of activation dynamics need further corroboration. Our findings add another puzzle piece in the dogs’ functional neuroanatomy and will be helpful in charting brain function across diverse species.

## Conflict of Interest

All authors declare no competing financial interests.

## Acknowledgements

This work would not be possible without the arduous work of our dog trainers, Marion Umek and Laura Lausegger. The authors would like to thank the student assistants in this project, Anna Thallinger, Olaf Borghi, and Sarah Binder. Finally, we thank the dogs and their owners for their patient and motivated participation. This project was supported by the Vienna Science and Technology Fund (WWTF) [10.47379/CS18012], the City of Vienna and ithuba Capital AG through Project CS18-012, the Austrian Science Fund, (FWF, Grant W1262-B29) and the Messerli Foundation (Sörenberg, Switzerland). The funders had no role in study design, data collection and analysis, decision to publish, or preparation of the manuscript.

## Author contributions

Conceptualization, design and methodology: CNAG, MB, RS, CL. Formal analysis and analysis plan: CNAG, CL. Plotting: CNAG, MB, LL. Writing manuscript draft: CNAG. Reviewing and editing drafts: CNAG, CL, MB, LL, SK, RS, LH. Feasibility testing: SK. Funding acquisition: LH, CL.

## Notes

https://osf.io/4gs9d/

### Competing Interest Statement

The authors have declared no competing interest.

